# Game Changing Mutation

**DOI:** 10.1101/2024.09.30.615800

**Authors:** Ziv Hellman, Omer Edhan

## Abstract

We present a model of the effect of mutation on haploid sexually reproducing populations by modelling the reproductive dynamics as occurring in the context of a common interests game played by the loci, with the alleles in the role of pure actions. Absent mutations, the population will deterministically converge to a pure Nash equilibrium of the game. A novel mutation adds new alleles, hence is tantamount to a change of the game by the addition of new actions. If the new game defined by the mutation removes the former pure Nash equilibrium the game changing mutation becomes in addition a Nash equilibrium changing mutation, as the population will then move to a new equilibrium with an increase in fitness. A graph of common interests games is defined, and evolution by mutation is modelled as a path through this graph.

**Short description:** Novel mutation in sexual reproduction evolution modelled as population shifts between Nash equilibria in common interest games

## 1 Introduction

The use of an evolutionary landscape, originally introduced in Wright (1931, 1932), has become part of the conceptual foundation of evolutionary biology. The landscape especially provides an appealing scaffolding for modelling the role of evolution in adaptive dynamics and hence is appears in many papers on the topic. In a typical presentation of the main concept, one models a population with a dominant genotypic type A, alongside variants of lower fitness that are small in population weight relative to the dominant type. Subsequently, a mutation of an allele in what had been a lower fitness variant B grants that type higher fitness value than A. This shifts the population to be mostly composed of genotypes of type B, in the process lifting the population higher up the landscape.

The literature on fitness landscape exploration by mutations is vast (see Kauffman and Levin (1987); Obolski et al (2018)). It works particularly well for studying adaptation by mutation in models of asexual reproductioion, where it is conceptually possible for a single allelic mutation to grant even a single individual in the population a fitness advantage that is directly transferred to descendants, enabling them to reproduce swiftly and take over the population.

In contrast, researchers have written about the difficulty of fitting models of sexual recombination to the adaptive landscape setting. They note that although on the one hand sexual recombination makes it likelier that beneficial mutations will become co-located in genotypes, on the other hand it can also undo beneficial combinations “because recently generated superior combinations are hard to maintain. They are likely to be lost because of recombination with other types” (Hadany and Becker (2003), see also Eshel and Feldman (1970)). The result of the sexual dynamics is thus typically presented in the literature as ambiguous, depending on the question of which force is stronger: the one pushing beneficial alleles together or the one tearing apart good combinations?

To contend with the difficulties of fitting sexual reproduction models to the landscape models, many models in the literature have added assumptions over and above the landscape model itself, such as assuming small population bottlenecks (Carson and Templeton (1984); Barton and Charlesworth (1984)), or specific heterogeneous distributions in the population (Hadany (2003); Weissman et al. (2010)).

We adopt a different and novel approach here to the study of mutation in sexually reproducing populations, moving away from the landscape, and modelling evolution as a path through an appropriately defined graph. We show that rather than an ambiguous tug-of-war between opposing forces, the sexual reproduction dynamics under full recombination leads asymptotically, in a determinstic trajectory depending on initial conditions, to a mononorphic population that is one of the Nash equilibria of an associated game. Mutations change the underlying game and possibly the equilibrium points as well, thus driving forward evolution.

Begin by noting that the main focus in the context of sexually reproducing populations is not on existing genotypes but rather on the collection 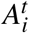 of alleles available at each locus *i* at time *t*, because the potential genotypes that may exist in the population will be composed of those alleles in various combinations. We call the profile 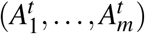 of available alleles, one per locus, an allelic formation. Regarding alleles as actions, we model sexual reproduction as a common interests game played by the loci: under pure strategies, each locus chooses an action/allele *a*_*i*_ from 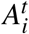, thus together forming a genotype *g* = (*a*_1_, *…, a*_*m*_). The payoff to forming genotype *g* is the fitness value of that genotype. Extending this to mixed strategies, with the probability of allele *a*_*i*_ being chosen equal to the prevalence of that allele in the population, yields the distribution of genotypes in the population at time *t*.

Against this background we can give the sexual reproduction dynamics a geometric interpretation. The space of possible genotypes, conditional on an allelic formation 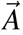, decomposes as a collection of basins of attraction in a space whose dimension is determined by the number of loci and the number of alleles per locus. The number of basins of attraction depends on the associated game 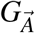, with one basin per pure Nash equilibrium of that game. Denoting by 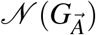 the set of pure Nash equilibria, under the sexual reproduction dynamics (without mutation), the population will move asymptotically towards one element of 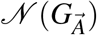, conditional on the initial starting point, in a trajectory of increasing fitness payoff until a pure Nash equilibrium is attained. In summary, sexual reproduction asymptotically yields a monomorphic population bearing the genotype of one of the Nash equilibria of a common interests game.

This leads to the following insight: a novel mutation added to the allele set of locus *i* at time *τ* is equivalent to the addition of a new action into the action set available to locus *i* as a player in a game. After such a mutation the loci are actually playing another game, which expands the game previously played by way of the addition of actions (thus also moving the dynamics into a space of higher dimension). We call such a mutation a game changing mutation.

Crucially, not every game changing mutation will have a long-term effect on the population; that depends on how the set of pure Nash equilibria changes under the expansion of the game. Denote by *G* the game played prior to the novel mutation and by *G*′ the game following the mutation; furthermore denote by ĝ ∈ *𝒩* (*G*) the Nash equilibrium genotype near which the pre-mutation population was located. If the geometry is such that ĝ is also a Nash equilibrium point in *𝒩* (*G*′), then post mutation the population will remain within a basin of attraction of *ĝ* (in the space of higher dimension), and asymptotically return towards ĝ, leading to no significant effect of the mutation.

In contrast, if *ĝ* is not a Nash equilibrium of *G*′, a significant change will be apparent in the population as it will move away from *ĝ* to a point *ĝ*′ in *𝒩* (*G*′). We term this event a Nash changing mutation. Under a Nash changing mutation, the fitness of the population increases as it moves towards a new Nash equilibrium point.

The model of mutations in sexually reproducing populations that emerges is of novel mutations expanding the sets of alleles available to populations. These mutations are all game changing in the sense that they change the underlying common interest game played by the loci, but not all such mutations move the population to a new equilibrium. Those that do so, the Nash changing mutations, will increase the population fitness in the process. In this picture, it is the Nash changing mutations that drive populations to new equilibria and higher fitness over time.

If we arrange the allelic formations in a directed graph, with an edge between two formations if and only if one is an expansion of the other by the addition of a single allele, evolution by mutations involves picking a path through this graph, via game changing or Nash changing mutations. Mutations that are not Nash changing have no long term effect on the mean population fitness, but the Nash changing mutations only increase the mean fitness. With probability one, the path evolution will take through the graph will be one of increasing mean fitness over time.

In addition, we show that this model can shed light on questions relating to fitness valley crossing, and evolutionary contingency, relating to the question of whether the order of mutations affects evolutionary outcomes.

In summary outline form, the main points of the model here are:

1. Under the sexual reproduction dynamics, absent mutations, formations of alleles may be considered to define a common interests game, where the players are the loci and their available actions are the alleles.
2. The game divides the space of genotypes into disjoint basins of attraction. Population dynamics follow trajectories of monotonically increasing population mean fitness within these basins until the population converges to a Nash equilibrium of the game, which is a fixed point.
3. Sparse non-novel mutations have negligible effects on the sexual reproductive trajectories, because they do not move the population from one basin to another.
4. Novel mutations are game changers: by adding new alleles they change the actions available to the players, hence changing the game
5. Not all game changing mutations, however, lead to population change. If the previous population fixed point, which was a Nash equilibrium of the previous game, is also a Nash equilibrium of the new game, the population will not move away from its previous equilibrium. One the other hand, if the previous equilibrium is not a Nash equilibrium of the new game, the population will move to a new equilibrium, in which case we term this event a Nash changing mutation.
6. With probability one, the path evolution will take is one of increasing mean fitness over time, from one game changing mutation to another at random arrival times.

## 2 Results

### 2.1 Overview of Model of Mutations

From an abstract perspective, loci, which represent sites at which alleles are located, are the most elementary aspect of the model. Each locus 1 ≤ *i* ≤ *m* is associated with a (possibly infinite) set *𝒜*_*i*_ of alleles. A *genotype* is a string of alleles, one at each locus. At any given time *t*, however, there is only a finite set 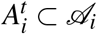 of alleles that are available at that time. The *m*-tuple 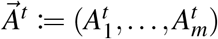 is termed the *allelic formation* at time *t*.

A population at time *t* is an ideal infinite collection of haploid genotypes, where each genotype *g* = (*a*_1_,, *a*_*m*_) satisfies the condition that for each 1 ≤ *i* ≤ *m*, allele 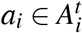 in other words, 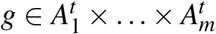 The collection of all genotypes present at time *t* when the allelic formation is 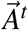 is denoted by 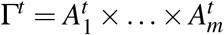. Each genotype *g* is associated with a fitness value *w*_*g*_ ∈ ℝ_+_, which is determined by *g* but in this model is independent of the population state and the time^1^. For simplicity we will assume that in each possible Γ there exists a unique genotype *g*^*^ ∈ Γ of maximal fitness.

Put together, we have all the ingredients for defining a common interests population game 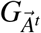 at time *t* from 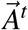, as follows. In this game, each locus 1 ≤ *i* ≤ *m* is a player. The actions available to player/locus *i* are the elements of the allele set 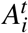. Each (pure) action profile *g* = (*a*_1_, …, *a*_*m*_) precisely corresponds to a genotype as defined above, and the payoff to player *i* when the action profile/genotype (*a*_1_, …, *a*_*m*_) is played is *w*_*g*_. This forms an elementary aspect of our model: reproduction modelled as a common interests game between loci.

The allelic formations can be arranged into a directed graph, with an edge from 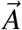 to 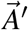 existing if and only if there is an 1 ≤ *i* ≤ *m* such that 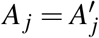 for every *j* ≠ *i*, but 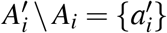 is a singleton (Figure 1). Since we have identified each allelic formation 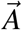 with a game 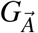 this graph can be perfectly mirrored in a corresponding graph of population games (Figure 2).

**Figure 1:**
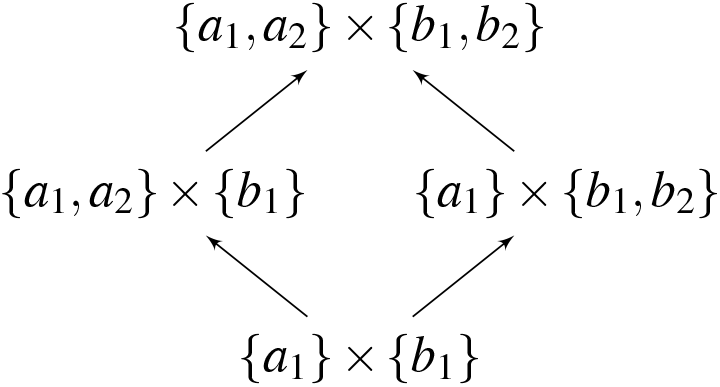
A schematic representation of an allelic graph.

**Figure 2:**
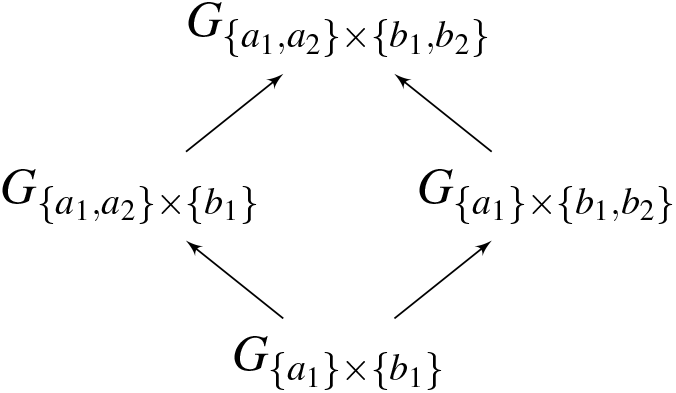
A schematic representation of a corresponding graph of population games.

Define the dimension of allelic formation 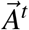 to be the dimension of the simplex Δ(Γ), where 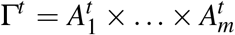. Define the dimension of the corresponding game 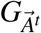 to be the same as the dimension of 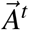. Then by construction the dimension increases as one moves up the graph of games (Figure 2).

A *novel mutation* is a single step in the graph of allelic formations in the direction of the arrows. A random walk through the graph yields a time-parametrised *novel mutation process* 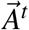, associating an allelic formation with each point in time *t*, from which one derives a corresponding time-parametrised process 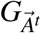 through the corresponding graph of games. The correspondence between the genotypic and game theoretic concepts is captured in Figure 3)

**Figure 3:**
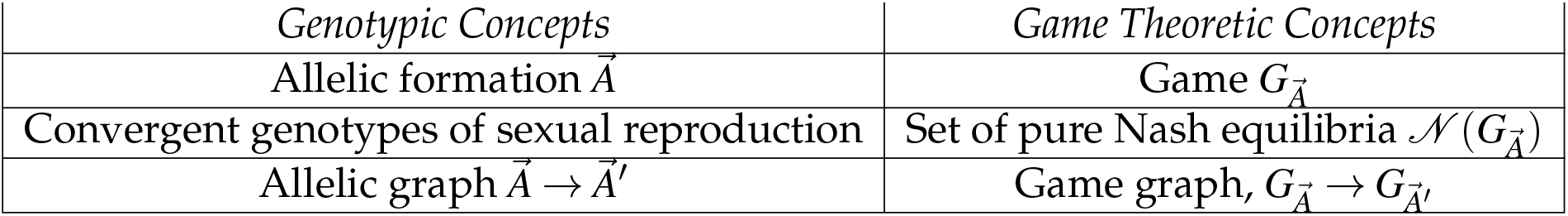
Parallels between genotypic concepts and game theoretic concepts in this paper

We note here a fact that will be important in the sequel: as one moves along a path up the population game graph, by construction all the actions available in games lower down the path are available in games farther up the path. However, this statement is not true for Nash equilibria: it is possible for a pair of games to be connected by an edge in the graph pointing from 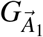 to 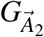 such that a profile that is a Nash equilibrium of 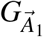 will not be a Nash equilibrium of 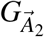, as the game geometry changes as dimensions are added. In other words the set of Nash equilibria of 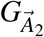 may not necessarily contain the set of Nash equilibria of 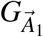.

Figure 4 provides a simple illustration of the general idea that adding dimensions may erase previous equilibria. On the left, a point marked *x* in a one dimensional simplex represents a stable equilibrium point, as forces on either side push any perturbation away from *x* back to the equilibrium. On the right, the addition of a dimension embeds this one dimensional simplex in a two dimensional simplex. The additional dimension enables trajectories that move away from *x*, hence in the two dimensional simplex *x* is not an equilibrium point.

**Figure 4:**
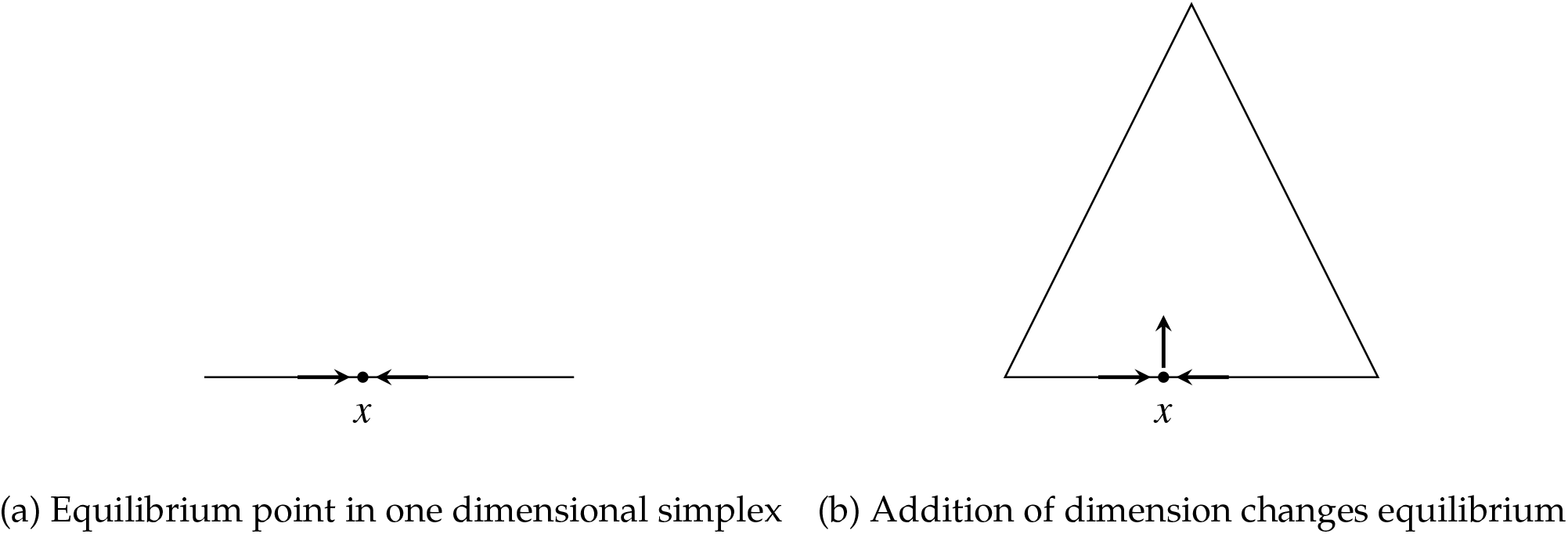
Illustration of added dimension removing equilibrium

### 2.2 Dynamics

A *game dynamic* is a map that assigns to each game 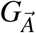 a differential equation 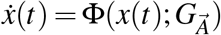, where *x* is a *population state* and 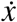 is the time derivative 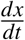 (see Sandholm (2010)). If the game 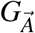 is not fixed but is instead given, e.g., by a novel mutation process 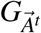, one obtains from this the dynamics 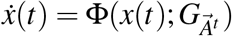.

If *τ* and *τ’* are the random arrival times of consecutive novel mutations, then the dynamic is determined by the same differential equation 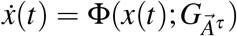 for all *τ* ≤ *t < τ’*. In this case, for *τ* ≤ *t < τ’*, the population state *x*^*t*^ will take values in the simplex Δ(Γ^*τ*^), whose dimension is fixed.

A random change then occurs at time *τ’*: the law of motion will become 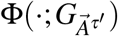 and hence the population state *x*^*t*^ will now take values in the higher dimensional simplex Δ(Γ^*τ’*^). The emerging dynamics is therefore a Piecewise Deterministic Markov Process (PDMP for short; see Davis (1994) and Benaïm et al. (2015)). However, whereas most PDMPs considered in applications take values in a space of fixed and bounded dimension (see Cloez et al. (2018)) this PDMP takes values in spaces of varying and perhaps unbounded dimension. The model does not preclude non-novel mutations; a non-novel mutation at any time *t* ∈ [*τ, τ’*) between two consecutive novel mutations in times *τ < τ’* can be captured by the deterministic dynamics of *x*^*t*^. For simplicity and coherence, we do not discuss such dynamics directly. However, some of our results show that the long run is mainly affected by novel mutations (see Theorem 4)

### 2.3 Asexual and Sexual Reproduction Dynamics

We are interested in two types of states: genotypic and allelic. *Genotypic states* are probability distributions *p*(*t*) ∈ Δ(Γ^*t*^). A genotypic state is *monomorphic* if there is a genotype *g* ∈ Γ such that *p*_*g*_(*t*) = 1. An *allelic state of locus i* is a probability distribution 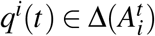. An *allelic state is* 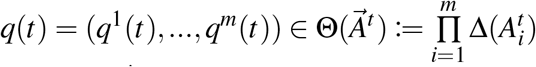 A genotypic state has an associated allelic state: denote by 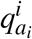 the proportion of genotypes *g* carrying allele *a*_*i*_ in locus *i*, i.e.,

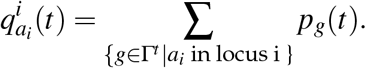

In models of asexual reproduction, genotypic states are typically the main focus of attention. The dynamics in such models is the replicator dynamics (Taylor and Jonker (1978)):

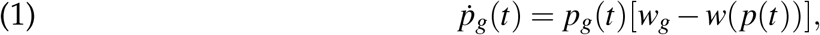

where 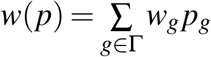 is the population mean fitness and the growth rate of *g* is *w*_*g*_ −*w*(*p*). As 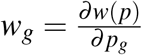, the mean fitness is a potential function of individual fitness and the population game in this case is a *potential game* (see Sandholm (2010)).

In models of sexual reproduction, allelic states are typically the main focus of attention. We assume that there is no correlation across alleles. In allelic state *q* the population mean fitness is given by 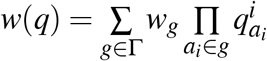, where *a*_*i*_ ∈ *g* means that allele *a*_*i*_ is in locus *i* of *g*. The individual payoff of player/locus *i* if she plays *a*_*i*_ (namely, allele *a*_*i*_ in locus *i*) is

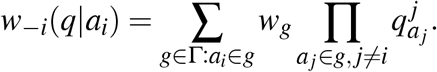

Similar to the above,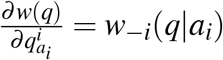, hence the mean fitness function *w* is a potential function for the game whose players are the loci.

If an allelic-formation 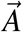 is kept fixed, then since the dynamics is defined via a potential function, it is natural to ask whether the dynamics is executing a gradient climb. For genotypic states (Equation (1)) it was shown by Shahshahani (1979) that when taking into consideration the appropriate metric^2^, this is indeed the case:

#### Theorem 1.

(Shahshahani (1979)). *The replicator dynamics of a potential game is a Shahshahani gradient in the interior of the simplex*.

From Shahshahani’s theorem it follows that in models of asexual reproduction without mutation, from any internal point in the simplex the population follows a trajectory of monotonically increasing mean fitness along the replicator dynamics, asymptotically approaching a monomorphic population comprised of the genotype of maximal fitness.

A similar result holds for sexually reproducing populations, as we show here. In the sexual reproduction model two individuals mate to produce offspring. When an individual with genotype *a* = (*a*_1_, *a*_2_, *…, a*_*m*_) mates with an individual with genotype *b* = (*b*_1_, *b*_2_, *…, b*_*m*_), the genotype of an offspring *c* = (*c*_1_, *c*_2_, *…, c*_*m*_) satisfies the following property: for each 1 ≤ *i* ≤ *m*, allele *c*_*i*_ equals either *a*_*i*_ or *b*_*i*_ with equal probability. The probability that an individual with genotype *g* will succesfully mate and produce offspring is proportional to *w*_*g*_; in other words, fitness is defined to be the probability of reproduction.

We denote by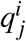 the proportion of gentoypes bearing the *j*-th allele in the *i*-th locus, and by 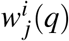 (*q*) the marginal fitness of that allele when the allelic state is *q*. Note that we are tracking here the proportions of alleles at each locus separately. Instead of one simplex to follow, there are *k* simplices, one simplex Δ_*i*_ for each locus *i*. Define the multi-simplex 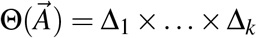 this is the state space of the sexual reproduction model.

Within each simplex Δ_*i*_ the alleles internal to locus *i* are competing with each other. However, the marginal fitness of each allele in locus *i* at each point in time is also a function of the full state of the population in the multi-simplex. The resulting *multi-replicator dynamics* is given^3^ by

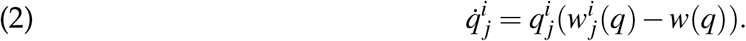

Despite the complexity of the sexual reproduction dynamic, even when the allelic formation 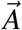 is kept fixed, it turns out that under an appropriate metric^4^ sexual reproduction executes a straight-forward gradient climb:

#### Theorem 2.

*For a fixed allelic formation* 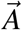, *the multi-replicator dynamics is a multi-Shahshahani gradient over a potential population game in the interior of the multi-simplex* 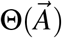.

#### Theorem 3.

*From any point in the interior of the multi-simplex, a sexually reproducing population with allelic formation* 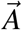 *converges asymptotically to a monomorphic population bearing a genotype that is in*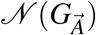, *the set of pure Nash equilibria of the associated common interests game*.

The exact definitions used in these theorems, and the proofs, can be found in the appendix. Intuitively, the common interests game divides the multi-simplex into disjoint basins of attraction, one basin of attraction for each pure Nash equilibrium in 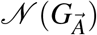. The population follows the multi-replicator dynamics, which determines gradient-climbing trajectories of monotonically increasing population mean fitness within each basin until the population converges to a Nash equilibrium of the game, which is a fixed point^5^.

There are immediate significant implications to these theorems: in asexually reproducing populations the mean fitness not only increases monotonically, but that it does so at a maximal possible rate of ascent. This mean fitness increase continues until the global maximum of the (in this case linear) mean fitness function over the convex simplex is attained, generically at a corner solution. Under this dynamics, for any *ε >* 0 the population will generically be *ε*-clustered around the genotype *g* of maximal fitness in finite time, and asymptotically will converge to *g*.

In sexually reproducing populations, the mean fitness also increases monotonically at a maximal rate of ascent, but now the mean fitness increase continues until the population asymptotically arrives at the local maximum of a basin attraction. Similar to the above, in finite time, for any *ε >* 0 the population will generically be *ε*-clustered around the genotype *g* of maximal fitness within one of the basins of attraction.

Chastain et al. (2014) also studies the relationship between games and sexual reproduction. Their model studies a discrete time algorithm, the multiplicative weights updating algorithm (MWUA), and shows that it corresponds to sexual reproduction under weak selection. The model there indicates that if the dynamics converges then sexual reproduction will achieve maximal fitness. However, the authors there do not show that the dynamics actually converges, nor that it constantly increases fitness. Although the MWUA is often called “the discrete replicator” it is important to note that in the no-regret regime studied in Chastain et al. (2014) the MWUA is not a discrete approximation of the replicator dynamics.

### 2.4 Game Changing and Nash Changing Mutations

In the model of this paper, relatively rare non-novel mutations have no significant effects on near equilbrium states of sexually reproducing populations. This is because such mutations can slightly change the distributions of the genotypes in the population, but if sufficiently rare they will not move the population out of one basin of attraction to another basin. As long as the population remains in the same basin of attraction, all the trajectories converge to the same pure Nash equilibrium of the associated game. We therefore concentrate from here on novel mutations.

In a slightly simplified version of the novel mutation process, we presume the existence of a Poisson clock with parameter *λ*. When the clock goes off at a random time *τ*, a locus *i* is selected at random together with 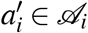, generating in this way a random walk through the allelic formations graph^6^.

Consider two separate times *τ*_1_ and *τ*_2_. We denote the the allelic formation at *τ*_1_ (resp., *τ*_2_) by 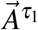 (resp.,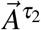), with associated game 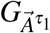 (resp.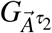). We also concentrate on genotypes *g*^1^ and *g*^2^ respectively where *g*^1^ is a Nash equilibrium of the game 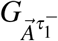 while *g*^2^ is a Nash equilibrium of the game 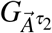

At time *τ*_1_, the allelic formation in the population is 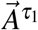 and the population is very nearly monomorphic at *g*^1^. Prior to *τ*_2_ no mutation occurs, and the allelic formation remains fixed, i.e., 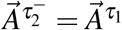. The population state remains near *g*^1^.

At random arrival time *τ*_2_ a mutation occurs, locus *i* is selected, and allele 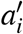 is added to 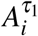, yielding 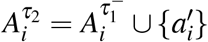. If 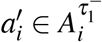, no change has occurred: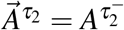, and the same population game as before continues to be played. In this case the population will not move away from its previous equilibrium point: *g*^2^ = *g*^1^.

Suppose instead that 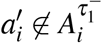. Then 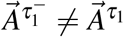, implying that a new game 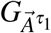 is being played. We term this a *game changing mutation* event.

This, however, does not necessarily mean that the population point moves to a new equilibrium point. The key question is whether *g*^1^, which is a Nash equilibrium of 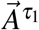is also a Nash equilibrium of 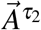. If yes, then the population will remain near *g*^1^ even after the game changing mutation. If no, since the population state which was a Nash equilibrium prior to the mutation is not a Nash equilibrium of the new game, the population will move towards the new equilibrium point at *g*^2^. In that case we say that a *Nash changing mutation* event has occurred.

This has implications for the population fitness level. If *g*^1^ = *g*^2^, the fitness level will remain at 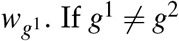 then 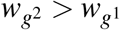 as the population moves to a new Nash equilibrium.

In summary, the following picture emerges. Evolution is modelled as a random connected path 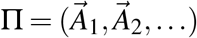 through the allelic graph.^7^ At each point *τ*_*j*_ in a series of random times (*τ*_1_, *τ*_2_, *…*), the population moves from 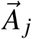 to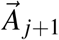.

If 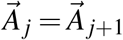, any mutation that has occurred is not a novel mutation. In this case, *g* ^*j*^ = *g* ^*j*+1^ and 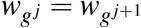 If 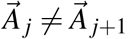 a game changing mutation has occurred. In this case there are two possibilities: either *g* ^*j*^ = *g* ^*j*+1^ and 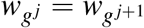, or *g* ^*j*^ ≠ *g* ^*j*+1^ and 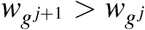. The latter event is a Nash changing mutation. As the population moves through the path Π, the mean population fitness generically increases monotonically.

This intuition is formalised in the following result (proved rigorously in the Appendix). This result has further implications for convergence to a monomorphic population, which is associated with pure Nash equilibrium of the underlying common interest game.

#### Theorem 4.

*Following a Nash changing mutation, the mean population fitness generically increases monotonically*.

#### Theorem 5.

*A population reproducing under the haploid sexual reproduction dynamics will with probability 1 follow a path through the allelic graph of (possibly weakly) monotonically increasing mean population fitness payoff*.

Theorem 5 summarises the main conclusion of our model: in haploid sexually reproducing populations, accumulating novel mutations that monotonically expand allelic formations lead to increasing fitness values.

Recall that the set of pure Nash equilibria of a game *G* is denoted *𝒢*. If novel mutations are rare, namely *λ* is sufficiently small, and if *τ < τ’* are two consecutive arrival times of novel mutations, one expects that with high probability in time *t* that is sufficiently near time *τ’*, the state *x*^*t*^ will be sufficiently near a Nash equilibrium of 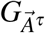. To capture this, denote by *d*(*x, 𝒩* (*G*)) the distance of a point *x* from the set of Nash equilibria of the game *G*. We then have the following result:

#### Theorem 6.

*Let τ < τ’ be two consecutive arrival times of novel mutations. For every ε >* 0 *there are δ >* 0 *and and a random variable λ*_0_ = *λ*_0_(*τ*) *such that λ*_0_ ∈ (0, ∞) *almost-surely and for almost-every realisation* 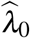 *we have that for every* 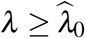 *and every t* ∈ [(1 −*δ*)*τ’, τ’*)

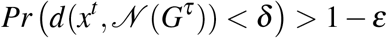

### 2.5 Fitness Valley Crossing

Raising fitness values may require combinations of mutations at different loci, since it is the interactions between alleles in the loci that determines fitness (examples from the literature include complex signalling pathways, and multiple mutations that may be needed to metabolise nutrients). It is possible that successive individually beneficial mutations can effect a monomorphic fitness climb, (Bridgham et al. 2006), but it is likelier that successful adaptations require combinations of mutations which individually are deleterious. When this is the case, it is said that populations need to cross a ‘fitness valley’ (Wright (1931)).

In this section we build on the haploid sexual reproduction model from the previous sections. Our results show that fitness valley crossing in our model may only require a single mutation, whereas common wisdom is that it requires at least two (e.g., Weissman et al. (2010) and Obolski et al. (2017)).

A fitness valley is given by two local maxima of the landscape *w*, say 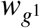 and 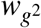 such that

a. 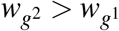
b. The Hamming distance *d*_*H*_ between the genotypes *g*^1^ and *g*^2^ is at least two, namely, at least two mutations are required to move from *g*^1^ to *g*^2^,
c. There is no local maximum *g*^3^ such that^8^ *d*_*H*_(*g*^1^, *g*^3^) + *d*_*H*_(*g*^2^, *g*^3^) = *d*_*H*_(*g*^1^, *g*^2^)

In the standard fitness landscape model of evolutionary biology the allele formation 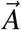 is kept fixed. Thus if *g*^1^ is a local maximum and the population state lies sufficiently near it then the population will converge to *g*^1^. In the event of mutation that changes a single allele of *g*^1^, the population state will be slightly perturbed initially, but will nevertheless continue to converge to *g*^1^; this is a corollary of Theorem 2. Thus a single mutation event that slightly changes the weights within the genotype space 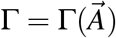 will not lead to a convergence to the improved maximum *g*^2^. This is referred to as the problem of *crossing the fitness valley*, the metaphor being of a landscape in a local fitness hill surrounded by a valley of lower fitness. As we have illustrated, in this model a population near a local fitness maximum will never climb to higher fitness values short of an extremely rare occurrence of two mutations within one individual.

Contrary to this, game changing mutations in a sexually reproducing population can dramatically and rapidly change the composition of a population. This can give sexually reproducing populations advantages over asexually reproducing populations.

#### Example 1.

Let *𝒜* = *{a*_1_, *a*_2_*}* and *ℬ* = *{b*_1_, *b*_2_, *b*_3_*}*. and let *A* = *{a*_1_, *a*_2_*}* = *𝒜* and *B* = *{b*_1_, *b*_2_*}* ⊂ *ℬ*.

Suppose that under initial conditions, Γ^*t*^ = *A × B* and Θ^*t*^ = Δ(*A*) *×* Δ(*B*). The fitness landscape, with fitness function *w*, is illustrated in Table 1 (with the Nash equilibria shown in bold), where *δ* is extremely small. It is captured graphically in Figure 5. This defines an allelic formation *C* = (*A, B*).

**Table 1:** Matrix 1.

| | $b_1$ | $b_2$ |
| --- | --- | --- |
| $a_1$ | **1** | $\delta$ |
| $a_2$ | $\delta$ | **2** |

**Figure 5:**
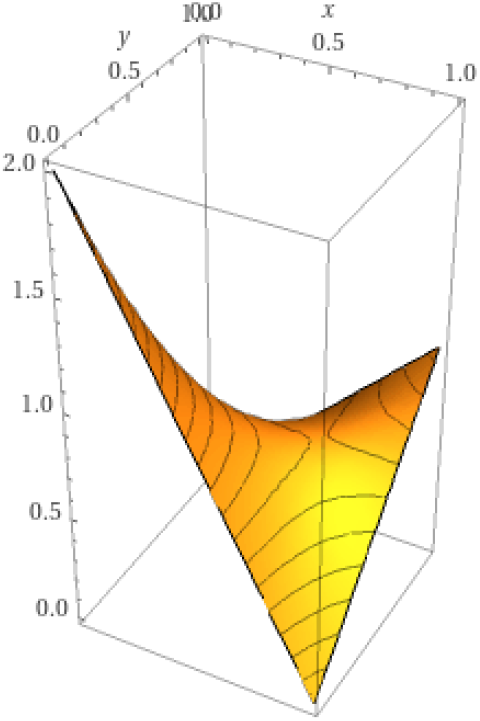
A plot of the game matrix 1 from Example 1, demonstrating a fitness valley.

Suppose that the initial population state *p*_1_ ∈ Δ(Γ^*t*^) places weight 1 − *ε* on genotype (*a*_1_, *b*_1_), and weight *ε/*2 on each of (*a*_2_, *b*_1_) and (*a*_1_, *b*_2_), with weight 0 on (*a*_2_, *b*_2_) (where *ε* is extremely small). In an asexually reproducing population, genotype (*a*_1_, *b*_1_) will maintain its central position in the population, leaving only trace amounts of individuals bearing genotypes (*a*_2_, *b*_1_) and (*a*_1_, *b*_2_). The mean population fitness value will be very close to one.

A sexually reproducing population will look very similar, apart from the fact that by random mating there will be small non-zero weight on genotype (*a*_2_, *b*_2_). A basin of attraction around genotype (*a*_1_, *b*_1_) will exist and the mean population fitness value will be very close to one.

Next, suppose that a mutation event occurs, with an individual bearing genotype (*a*_1_, *b*_1_) mutating to (*a*_1_, *b*_3_). The fitness landscape then changes to that illustrated in Table 2.

**Table 2:** Matrix 2.

| | $b_1$ | $b_2$ | $b_3$ |
| --- | --- | --- | --- |
| $a_1$ | 1 | $\delta$ | $\frac{3}{2}$ |
| $a_2$ | $\delta$ | **2** | $\frac{7}{4}$ |

For an asexually reproducing population, this will have the effect of changing the state of the population to being nearly monormophically (*a*_1_, *b*_3_), and the mean population fitness value will be nearly 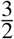.

The result in the sexually reproducing population will be quite different. The allelic formation expands to *C*′ = (*𝒜, ℬ*). As Table 2 indicates, (*a*_1_, *b*_1_) is not a Nash equilibrium of the expanded game. Since the dynamics must carry the population towards a Nash equilibrium, the population will move away from (*a*_1_, *b*_1_) and will eventually be nearly monomorphically composed of genotype (*a*_2_, *b*_2_). The mean population fitness value will be nearly 2.

It is interesting to note that in this example although the mutation to allele *b*_3_ kicked off the process, its appearance eventually boosts allele *b*_2_ asymptotically to fixation while *b*_3_ itself asymptotically goes extinct, as illustrated in Figure 6. An observer who views the population when it is mainly composed of genotype (*a*_1_, *b*_1_) and later views it when it is mostly (*a*_2_, *b*_2_) may not have an indication that a mutation to (*a*_1_, *b*_3_) was involved at all. ♦

**Figure 6:**
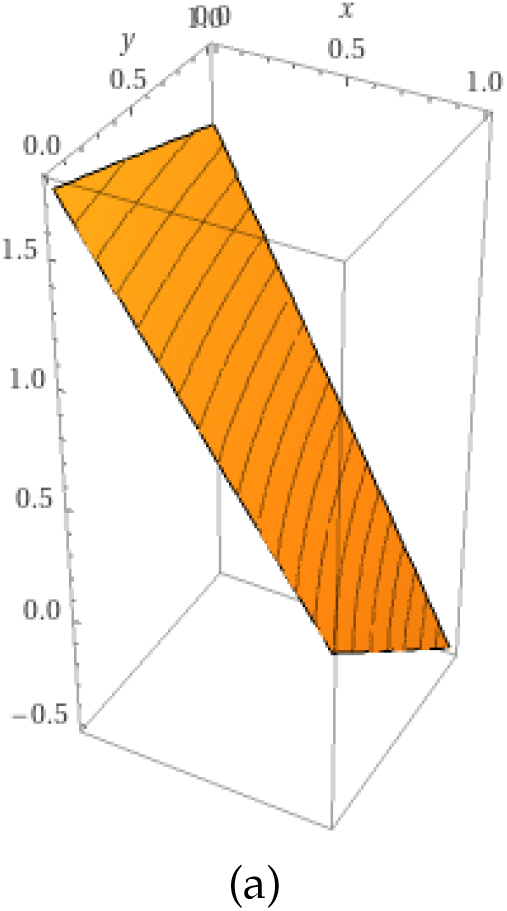
A plot of a section of game 2 in which the probabilities are equal across the diagonal, namely *x*(*b*_1_) = *x*(*b*_2_). Here, the valley of Figure 5 is replaced by a steep climb.

The previous example exhibited a situation in which sexual reproduction gains an advantage over asexual reproduction by way of game changing mutations. The sexual reproduction dynamics, however, can also be disadvantageous in certain situations. Its main weakness is that a mutation that does not remove a Nash equilibrium around which a population is clustered will not move the population to a new state, even if that mutation is highly beneficial for the genotype in which the mutation has occurred. The next example exhibits this.

#### Example 2.

This example is very similar in its initial conditions to Example 1. Let *𝒜* = *{a*_1_, *a*_2_*}* and *ℬ* = *{b*_1_, *b*_2_*}*, Γ^*t*^ = *𝒜 ×ℬ*. The fitness landscape is again that of Table 1.

Suppose again that the initial population state *p*_1_ ∈ Δ(Γ^*t*^) places weight 1 −*ε* on genotype (*a*_1_, *b*_1_), and weight *ε/*2 on each of (*a*_2_, *b*_1_) and (*a*_1_, *b*_2_), with weight 0 on (*a*_2_, *b*_2_) (where *ε* is extremely small). As before, both the asexual and sexual populations are clustered around genotype (*a*_1_, *b*_1_) with mean population fitness near one (in the sexually reproducing population, however, there will rapidly be small population weight developing on genotype (*a*_2_, *b*_2_), via recombination, as opposed to the zero weight on (*a*_2_, *b*_2_) of the asexual population).

Suppose now that an individual in the population bearing genotype (*a*_2_, *b*_1_) undergoes a mutation to (*a*_2_, *b*_2_). In the sexually reproducing population, nothing changes: there was already a small weight of individuals with genotype (*a*_2_, *b*_2_), hence the mean population fitness remains near one. In the asexual population, however, this mutation brings about a dramatic change: the introduction of an individual of genotype (*a*_2_, *b*_2_) leads asymptotically to that genotype taking over the population, driving the mean population fitness towards 2. ♦

### 2.6 Evolutionary Contingency

Both asexual and sexual reproduction strive to find maximal-fitness solutions. However, this process may not be deterministic as the outcomes may also depend on idiosyncratic events that an evolving lineage experiences such as the order of appearance of random mutations. Should the tape of life be replayed, would it produce similar living beings? This question known as *historical contingency*, or *contingency* for short, was argued by Stephen Jay Gould (Gould (1989)) to be an essential feature of evolution.

Gould’s original idea introduced confusion regarding the notion of contingency and the way it operates (Blount et al. (2018)). Some authors have tried to resolve this confusion (e.g., Beaty (2011), Beaty (2017), Blount et al. (2018), Desjardins (2009), and Desjardins (2011)), but these papers generally did not present thorough modeling frameworks.

In the standard models of asexual mutation in the literature, the order in which a chain of mutations occurs makes no difference to the end resulting genotype (although it could affect the chances that a particular chain will arrive at the endpoint). Consider, for example, two chains of mutations 1) one taking the population from monomorphic (*a, b*) to (*c, b*) and then to (*c, d*); 2) alternatively another path of genotypes moving the population from (*a, b*) to (*a, d*) and then to (*c, d*). In both cases, the end result is (*c, d*) and the payoff is that of the genotype (*c, d*).

In the model of haploid sexual reproduction of this paper, however, the end result of a chain of mutations may be very dependent on the order in which mutations occur, as the population may climb the genotypic lattice through different paths from one point to another. This is exhibited in the following example.

#### Example 3.

Recall the matrices in Tables 1 and 2 above. We have already established that when Table 1 is augmented to Table 2 the sexually reproducing population will move, from a state close to (*a*_1_, *b*_1_), to (*a*_2_, *b*_2_).

Consider now the possibility of Matrix 1 being augmented instead to Matrix 3 (see Table 3). In the move from Matrix 1 to Matrix 3, the state (*a*_1_, *b*_1_) is no longer a Nash equilibrium. The population will instead move to the state (*a*_3_, *b*_1_).

**Table 3:** Matrix 3.

| | $b_1$ | $b_2$ |
| --- | --- | --- |
| $a_1$ | 1 | $\delta$ |
| $a_2$ | $\delta$ | 2 |
| $a_3$ | 3 | $\delta$ |

Finally, consider Matrix 4 (see Table 4). If Matrix 2 is augmented to Matrix 4, from a state close to the (*a*_2_, *b*_2_), the population will remain in the vicinity of (*a*_2_, *b*_2_). Similarly, if Matrix 3 is augmented to Matrix 4, from a state close to the (*a*_3_, *b*_1_), the population will remain in the vicinity of (*a*_3_, *b*_1_).

**Table 4:** Matrix 4.

| | $b_1$ | $b_2$ | $b_3$ |
| --- | --- | --- | --- |
| $a_1$ | 1 | $\delta$ | $\frac{3}{2}$ |
| $a_2$ | $\delta$ | 2 | $\frac{3}{2}$ |
| $a_3$ | 3 | $\delta$ | $\delta$ |

Hence we have shown the path dependence of the game changing mutation process: adding *b*_3_ and the *a*_3_ does not lead to the same result as adding *a*_3_ first and then *b*_3_. The former represents moving from Matrix 1 to Matrix 2 to Matrix 4, hence from (*a*_1_, *b*_1_) to (*a*_2_, *b*_2_), while the latter moving from Matrix 1 to Matrix 3 to Matrix 4, hence from (*a*_1_, *b*_1_) to (*a*_3_, *b*_1_). ♦

### 2.7 Robustness under Random Arrivals

A powerful recasting of evolutionary contingency is the predictability of evolutionary outcomes (Orgogozo (2015)): if life’s tape is replayed can we make predictions about what to expect? In the next two section we explore two matters related to predictability. The first, which we discuss here, is the robustness of the order of arrival of novel mutations to randomness.

Our previous examples concentrated on realised arrival orders of mutations, illustrating the underlying idea of game changing and Nash changing mutations. Here we extend the discussion to accommodate random arrivals of mutations and evaluate the probability of different contingencies.

We revisit Example 3, by assuming that the arrivals of mutations for actions *a*_3_ and *b*_3_ follow exponential distributions with parameters *λ*_*a*_ and *λ*_*b*_ respectively. Let *τ*_*a*_ and *τ*_*b*_ denote the random arrival time of actions *a*_3_ and *b*_3_ respectively. As we have seen, the resulting Nash equilibrium for the event *τ*_*a*_ *< τ*_*b*_ will be different from that of the event *τ*_*b*_ *< τ*_*a*_. To evaluate the probability of each Nash equilibrium, one needs to evaluate the probability of the aforementioned inequalities between arrival times:

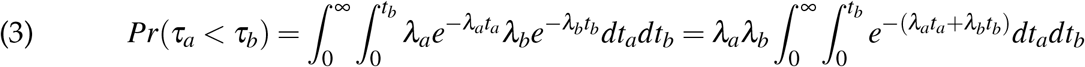

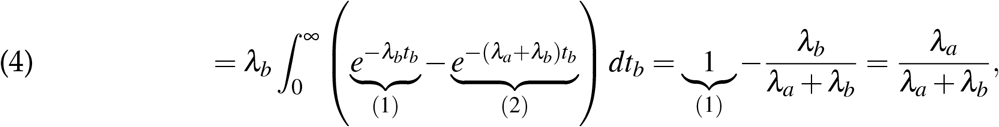

and by symmetry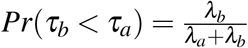.

It is easy to seen that, e.g., as 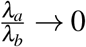 we have *Pr*(*τ*_*b*_ *< τ*_*a*_) → 1, namely, as the arrival time of *b*_3_ becomes large compared to that of *a*_3_, the probability of the Nash equilibrium moving from (*a*_1_, *b*_1_) to (*a*_3_, *b*_3_) tends to 1, showing the robustness of Example 3 in this case.

### 2.8 Replaying the Tape of Life

What does assessing the ‘predictability’ of outcomes mean? Example 3 showed how differences in fitness emerge from different orders of mutations. Here we aim at finding a quantitative estimation of this effect. As we shall see, the measurement of fitness isn’t the only important thing - measurement timing is important as well.

Let 0 = *τ*_0_ *< τ*_1_ *< τ*_2_ *<* … *< τ*_*k*_ *<* be the random arrival times of novel mutations, and define

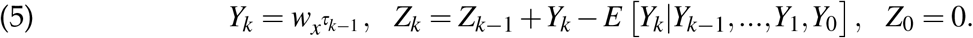

Hence *Z*_*k*_ defines the random walk of fitness due to novel mutations. As *Z*_*_ is a martingale, we could apply the martingale CLT to study it. Suppose that fitness jumps are bounded, namely there is a *c >* 0 such that |*Z*_*k*+1_ −*Z*_*k*_| *< c* for every *k*. Define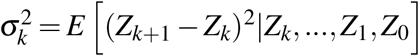 and let

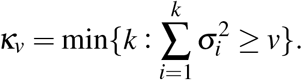

Then

#### Theorem 7.

*The random variable* 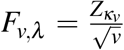*converges in distribution to the normal distribution with mean* 0 *and variance* 1 *as v* → ∞.

Notice that the measurement timing is important for the result. The measurement time *κ*_*v*_ itself is a random variable. Furthermore, the process contains randomness from two different sources:

1. The random order of arrival of novel mutations; and
2. The distance of the state *x*^*t*^ from the respective pure Nash equilibrium.

As *λ* → 0 one would expect the randomness due to item (2) to diminish, hence we would be left with the randomness due to random order of arrival of random mutations.

## 3 Conclusion and Further Questions

We have introduced a model of mutation as stochastic movement through the allelic lattice under the haploid sexual reproduction dynamic. This leaves us with several questions for further research:

1. The model here presumes that novel alleles are always added to existing allelic formations; alleles never disappear. This is unrealistic. A model in which alleles can appear over time and also disappear is needed. In such a model, paths through the allelic graph will not be unidirectional, and at present it is unclear what long term results with respect to mean population fitness values could be expected; further assumptions to the model may be necessary.
2. What qualitative long term convergence results can be expected in the model of this paper? Benaïm et al. (2015) developed such a theory for certain piecewise deterministic Markov processes (PDMP) with bounded dimension. Our work here introduces a new example of PDMP whose ‘dimension’ (measured here by the size of the allelic formation) may be unbounded as *𝒜*_*i*_ may be infinite. Developing such a theory for PDMPs similar to the one we have discussed is beyond the scope of this paper, and will be pursued in a subsequent work.
3. How likely are new Nash equilibria to appear as games are expanded? This is a question that is of interest in game theory in general, not only with respect to evolutionary theory: as games expand, as defined in this paper, should one expect new Nash equilibria to apepar often or rarely? The answer may depend on specific structural aspects of the games involved. A characterisation of such aspects would be a contribuction to the literature.
4. How much of what is presented here survives in models with frequency dependent fitness, or in models of finite populations? A model with frequency dependence will likely not maintain the the structure of a strategic game on which many of the results here depend – the less demanding concept of a population game will be relevant. In a finite population model, genetic drift effects may become prominent to the point that they significantly change the convergence results presented in the body of this paper.

## Appendices

### A Appendix

#### A.1 Derivation of the Sexual Multi-Replicator Equation

Asexual reproduction manifests replicator equation dynamics, which enables one to make use of results regarding the replicator equation to study asexual reproduction. We shown in this section that the multi-replicator can analogously be used to study sexual reproduction dynamics (in the mutation-free model) in continuous time.

Recall that we have denoted the fitness of a genotype *g* by *w*_*g*_, which we naturally interpret here as the instantaneous growth rate of the share of the mass of the population bearing genotype *g*.

When we adopt the perspective of the loci in the sexual reproduction model as playing a game, if *g* = (*a*_1_, *…, a*_*m*_) is regarded as the pure action profile of *m* loci, we can denote by *w*^*i*^(*g*) the payoff to locus *i* when action profile *g* is played. Since this game is a common interests game whose payoff is the fitness of *g*, we have *w*^*i*^(*g*) = *w*_*g*_ for all *i*.

Let *A*_*i*_ denote the set of alleles of locus *i* at time *t* (where for clarity we suppress the explicit expression of *t* from here on). Let 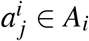 denote a particular allele in *A*_*i*_. Denote 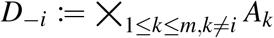 Let 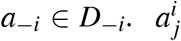 and *a*_−*i*_ together define a genotype 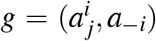. We will term *a*_−*i*_ the partners of 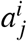 in forming *g*. We may then conceive of the set *D*_−1_ as representing all possible partners of 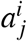 for any *j* with respect to the set of all possible genotypes containing 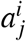.

Moving from pure to mixed strategies, denote a mixed strategy of player *i* in the common interests game by 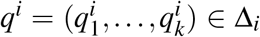. Here 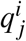 denoted the share of the population with allele *j* at locus *i* (hence by definition 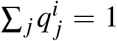). A profile of mixed strategies is a point *q* = (*q*^1^, *…, q*^*m*^) ∈ Θ. An element of ∏_1≤ *ℓ* ≤*m*; *ℓ*≠*i*_ Δ(*A*_*ℓ*_) is denoted *q*_−*i*_.

Furthermore, denote by *p* the total mass of the population at time *t*, and by 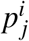 the mass of the population bearing allele *j* of locus *i*. What we have denoted by 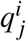, the share of the population with allele *j* at locus *i*, is then given as

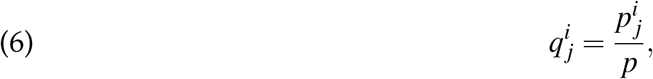

equivalently, 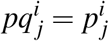.

The joint application of a joint strategy profile *q* by all the loci yields an expected payoff for locus *i* that we will denote *w*^*i*^(*q*), extending the notation *w*^*i*^(*g*) to mixed strategy profiles. In fact, since the game being played is a common interests game, with each locus receiving the same payoff for each pure strategy, the mean population fitness under mixed strategy *q* is equal to the expected payoff *w*^*i*^(*q*) for each single locus *i*. In other words, denoting the mean population fitness under *q* by *w*(*q*)

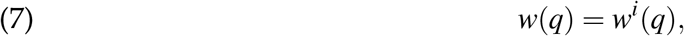

for any locus *i*.

Given 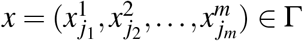 ∈ Γ and a profile of mixed strategies *q*, denote

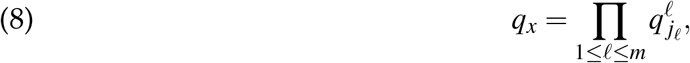

and

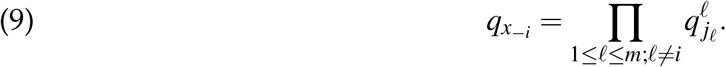

An interpretation of Equation (9) is that it expresses the probability of 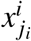 encountering the partners *x*_−*i*_ under the mixed strategy *q*.

Next we want to specify the payoff that locus *i* receives for placing weight 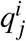 on allele 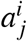 when the overall expected payoff to locus *i* is *w*^*i*^(*q*).

Denote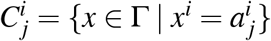, i.e., this is the set of all possible genotypes when the allele of locus *i* is fixed at 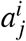 and all possible partners are joined to it. Suppose that those possible partners are distributed according to *q*_−*i*_. In that case, the *marginal fitness* of allele 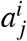 (with respect to mixed strategy profile *q*) is

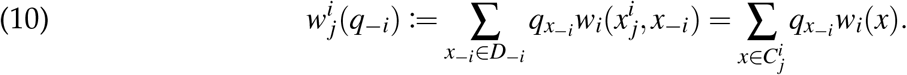

Equation (10) expresses the marginal fitness of allele 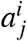 under *q* as the weighted average of the payoff to locus *i* under the genotypes *x* containing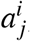, with the weightings given by *q* ._*x*−1_

Finally, note that

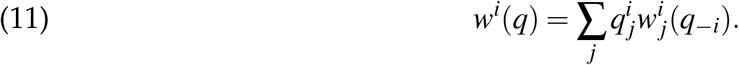

Recalling that *w*^*i*^(*q*) = *w*(*q*) for each *i* (by Equation (7), essentially expressing the fact that the game is a common interests game), we can calculate, for each *i* and *j*:

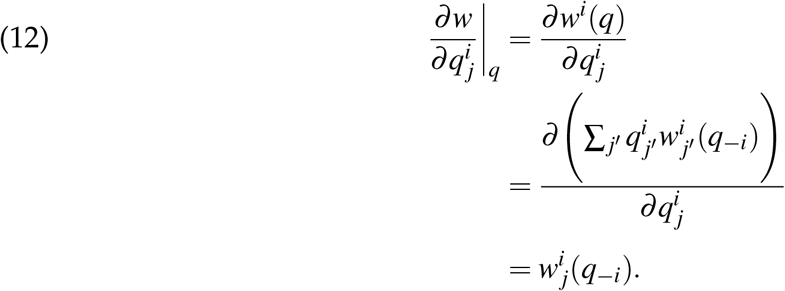

From the perspective of any single locus *i*, each allele 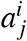 is competing for a frequency share in the population against the other alleles in the same locus. At a state of the population *q* ∈ Θ, the payoff to allele 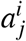 is the marginal fitness 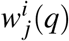. The greater 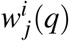, the faster the share of 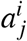 grows. This hints that the alleles at each locus are undergoing a replicator-like dynamic. This can be shown formally.

From Equation (6), one attains 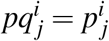. Differentiating this with respect to *t* yields 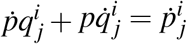. Moving terms around,

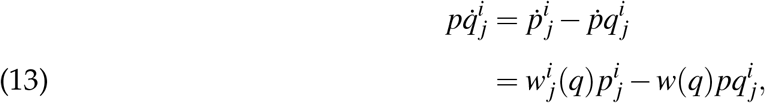

where we have used the relations 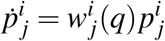 and 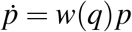 which hold by the definitions of 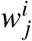 and *w* as instantaneous rates of growth.

Dividing both sides of Equation (13) by *p* gives

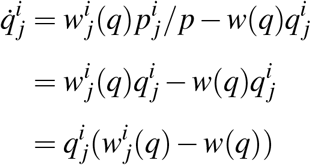

or more succinctly

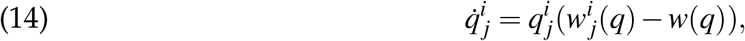

for each *i* and *j*, which is what we have called the multi-replicator equation.

#### A.2 Geometry Preliminaries

Recall that a main construct of interest throughout this paper is a polytope composed of an *m*-cross product of simplices, where each simplex Δ_*i*_ is a *k*-simplex with vertices over *A*_*i*_:

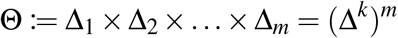

where Δ_*i*_ is understood as short-hand for Δ(*A*_*i*_). We have called Θ thus defined a multi-simplex. Each simplex Δ_*i*_ is a subset of ℝ^*k*+1^. Hence Θ ⊂ ℝ^*k*+*m*+1^.

We adopt the following conventions for denoting elements of a multi-simplex Θ. Firstly, we suppose that the constituent simplices (Δ_1_, *…*, Δ_*m*_) are ordered from 1 to *m*, and furthermore that within each simplex Δ_*i*_ the vertices of the simplex 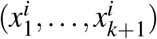 are similarly ordered. In general, 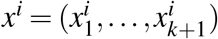 will denote an element of Δ_*i*_, and 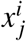 will denote the *j*-th element of *x*^*i*^. Occasionally we will wish to order all of the vertices of all of the constituent simplices Θ together; in that case we will write *y* ∈ Θ as *y* = (*y*_1_, *…, y*_*m*(*k*+1)_).

A vector *v* ∈ ℝ^*k*+1^ is tangent to Δ^*k*^ at a point *p* if and only if the total derivative *Dψ*_*p*_(*v*) = 0, i.e.,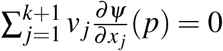. This translates into the condition Σ _*j*_ *v* _*j*_ = 0.

With regards to the multi-simplex Θ = (Δ^*k*^)^*m*^, first denote by **1**^*m*^ the *m* product of the *k* + 1 vector **1** defined above, and then define the mapping

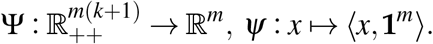

Then (Δ^*k*^)^*m*^ = (Ψ)^−1^(1). A vector *v* ∈ ℝ^*m*(*k*+1)^ is tangent to Θ at a point *p* iff for each 1 ≤ *i* ≤ *m*, one has 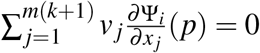. This translates into the condition Σ *v* _*j*_ = 0 separately for each *i*.

A Riemannian metric *h* over a manifold *M* associates a symmetric, positive-definite matrix *g*(*x*) = (*g*_*ij*_(*x*)) to each point *x* ∈ *M* in a smooth manner.

Denote the tangent space to *M* at a point *x* by *T*_*x*_*M* and denote the Euclidean inner product of two vectors *ζ, η* in *T*_*x*_*M* by *(ζ, η)*_*x*_. Then for a Riemannian metric *h* over a manifold *M* the inner product of *ζ, η* at *x* with respect to *h* is

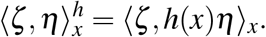

If *F* is a real-valued smooth function over manifold *M* with a Riemannian metric *h* then the derivative *DF*(*x*) is a linear map from *T*_*x*_*M* to ℝ. A vector *η* ∈ *T*_*x*_*M* such that

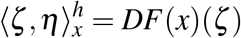

for all *ζ* in the tangent space at *x* is called the *gradient* with respect to the metric *h* at *x* (Hofbauer and Sigmund (1998)). Such a gradient is conventionally denoted ∇_*h*_*F*(*x*). A smooth vector field 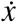 determining a dynamical system is an *h-gradient field* if 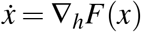 for all *x* ∈ *M*.

The Shahshahani metric, restricted to the open interior of a *k*-simplex, is defined by the metric tensor

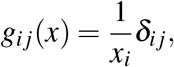

for *x* ∈ Δ^*k*^ and 1 ≤ *i, j* ≤ *k* + 1; equivalently, this is the (*k* + 1) *×* (*k* + 1) diagonal matrix whose non-zero entries are 1*/x*_*i*_. The simplex together with this metric is called the Shahshahani manifold. A gradient field with respect to the Shahshahani metric is called a Shahshahani gradient.

When we focus not on one simplex but rather the multi-simplex Θ := Δ_1_ *×* Δ_2_ *×*… *×* Δ_*m*_ = (Δ^*k*^)^*m*^, the Shahshahani metric tensor is insufficient. We therefore extend this to a metric tensor, which we call the *multi-Shahshahani metric*, over the product (Δ^*k*^)^*m*^ by defining

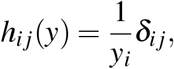

for *y* = (*y*_1_, *…, y*_*m*(*k*+1)_) ∈ (Δ^*k*^)^*m*^, and 1 ≤ *i, j* ≤ *m*(*k* + 1). This looks superficially identical to the Shahshahani metric but it should be kept in mind that here 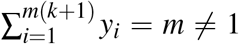 only the elements restricted to each of the individual simplices comprising Θ sum to 1. This results in quite a different manifold. One way to understand the multi-Shahshahani metric is that if it is projected on to any single constituent simplex Δ_*i*_ one recapitulates the standard Shahshahani metric over Δ_*i*_.

It is well known that the replicator dynamics is a gradient flow along the Shahshahani metric. When a population following the replicator dynamics is far from a fitness peak, the gradient climb is relatively steep; the nearer the population is to the peak, the less steep the gradients become.

The multi-replicator dynamics is a gradient flow according to the multi-Shahshahani metric, but at the same time it executes a replicator dynamics internally at each locus. In a sense the multi-replicator is a more ‘modular’ approach to evolution than the replicator: it simultaneously enables gradient ascents of differing slopes in the separate loci, in contrast to the replicator dynamics, which takes into account entire genotypes, not loci.

#### A.3 Proof of Theorem 2

We present here a proof of Theorem 2, which we restate:

For a fixed allelic formation 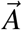, the multi-replicator dynamics is a multi-Shahshahani gradient over a potential population game in the interior of the multi-simplex Θ 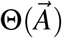.

From any point in the interior of the multi-simplex, a sexually reproducing population converges asymptotically to a monomorphic population bearing a genotype that is in 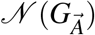, the set of pure Nash equilibrium of the associated common interests game.

*Proof*. Let Θ = Δ_1_ *× … ×* Δ_*m*_, and let *x* = (*x*^1^, *…, x*^*m*^) ∈ Θ. Let 1 ≤ *i* ≤ *m* be arbitrary, and let *ζ*^*i*^ be a vector in the tangent space to Δ_*i*_ at *x*^*i*^. Let *h*_*i*_ be the projection of the multi-Shahshahani metric *h* to Δ_*i*_. Then, since *h*_*i*_ is a Shahshahani metric with respect to Δ_*i*_,

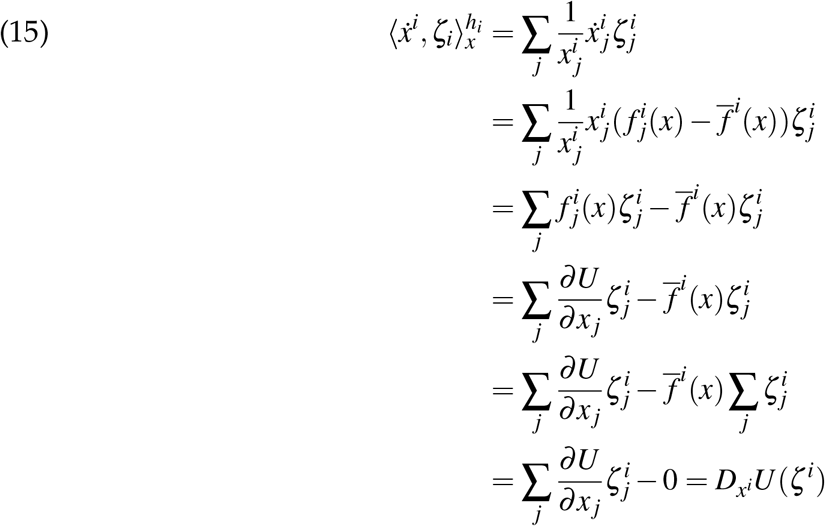

showing that 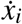 is a gradient with respect to *h*_*i*_.

Up to here, the focus was on the perspective of a particular *i*. Making use now of the assumption of the tensor *h* over Θ as being composed as *h*(*x*) = (*h*_1_(*x*^1^), *h*_2_(*x*^2^), *…, h*_*m*_(*x*^*m*^)), the decomposition of any tangent vector *ζ* with respect to Θ as *ζ* = (*ζ*_1_, *…, ζ*_*m*_) yields

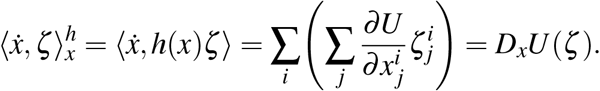

We conclude from this that the law of motion of the multi-replicator induces a multi-Shahshahani gradient vector flow.

#### A.4 Proof of Theorem 3

We present here a proof of Theorem 3, which we restate:

From any point in the interior of the multi-simplex, a sexually reproducing population with allelic formation 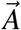 converges asymptotically to a monomorphic population bearing a genotype that is in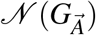, the set of pure Nash equilibrium of the associated common interests game.

*Proof*. We have already established elsewhere in this paper that under the sexual replicator dynamics, the game 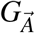 can be interpreted as a common interests game following a multi-replicator dynamics in the multi-simplex, which by Theorem 2 induces a multi-Shahahani gradient vector flow. The population will therefore follow a trajectory of increasing mean fitness, which can only end, asymptotically, at a point of local fitness maximum, which in game theoretic terms will be a pure Nash equilibrium in the set 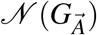.

#### A.5 Proof of Theorem 4

Recall the statement of the theorem:

Under a Nash changing mutation, as a population moves from near-monomorphically bearing genotype *g*^*t*^ to genotype 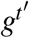, the mean population fitness generically increases monotonically.

*Proof*. By definition, when a game changing mutation occurs, genotype *g*^*t*^, which had been a Nash equilibrium of the previous game *G*_*B*_, is not a Nash equilibrium of the post-mutation game *G*_*B’*_. Therefore, immediately after the mutation, the population is near a point that is not a Nash equilibrium. It will then commence immediately to climb the multi-Shahshahani gradient over *G*_*B’*_, increasing the mean population fitness monotonically, until it attains a local maximal point near a Nash equilibrium point of the game *G*_*B’*_ represented by genotype 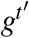.

#### A.6 Proof of Theorem 5

Recall the statement of the theorem:

A population reproducing under the haploid sexual reproduction dynamics will with probability 1 follow a path through the allelic graph of (perhaps weakly) monotonically increasing mean population fitness payoff.

*Proof*. The random process of mutations of the main body of the paper will with probability 1 yield a sequence of game changing mutations, some of which will also be Nash changing.

A Nash changing mutation increases population fitness, as the population moves to a new equilbrium point. A game changing mutation that is not Nash changing returns the population back to the Nash equilibrium point at which it was prior to the mutation, with no change in fitness payoff. In between mutations, the population remains near a Nash equilbrium. It follows that with probability 1 the dynamics follows a path of at least weakly monotonically increasing fitness payoff.

#### A.7 Proof of Theorem 6

Recall that the Theorem states that if *τ < τ’* are two consecutive arrival times of novel mutations, then for every *ε >* 0 there are *δ >* 0 and and a random variable *λ*_0_ = *λ*_0_(*τ*) s.t. *λ*_0_ ∈ (0, ∞) almost-surely and for almost-every realisation 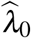 we have that for every 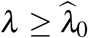 and every *t* ∈ [(1 −*δ*)*τ’, τ’*)

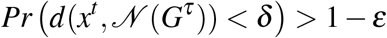

To prove this, let *x*^*^ be the random equilibrium s.t. without any further novel mutation the dynamics, when initiated at *x*^*τ*^, converges to *x*^*^. Let *B*_*δ*_ = *{y* : ||*y* − *x*^*^|| *< δ}*. There is a random time *T* = *T* (*x*^*τ*^, *δ*) s.t. if 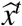 is the dynamics initiated at *x*^*τ*^ without any further novel mutation then for every time *t > T* we have 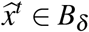 ∈ *B*_*δ*_. To finish the proof choose *λ*_0_ = *λ*_0_(*τ*) s.t. *Pr*((1 −*δ*)*τ’* −*τ > T*) *>* 1 − *ε*.

#### A.8 Proof of Theorem 7

The theorem is a restatement of the martingale Central Limit Theorem for this case (Hall and Heyde (1980)).

## Footnotes

1 This explicitly does not exclude the possibility of epistasis between loci.

2 Known as the *Shahshahani metric*, see Appendix A.2

3 See Appendix A.1 for a full derivation of the dynamics.

4 Which we call the *multi Shahshahani metric*, see Appendix A.2 for further details.

5 For discrete time models, results very similar to those in this section can be found in Palaiopanos et al. (2017), Novak and Barton (2017), and Edhan et al. (2021). The theorems here extend those results to continuous time models.

6 If 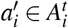, no walk subsequently occurs in the graph.

7 Since the allelic graph was constructed as a directed graph, with each edge an arrow whose end point is an allelic formation containing more alleles than the allelic formation at the starting point, alleles are only added along the path Π, never removed. In a more general model one may consider paths that also move ‘backwards’ along the edges, which would correspond to loss of alleles.

8 The triangle inequality *d*_*H*_(*g*^1^, *g*^3^) + *d*_*H*_(*g*^2^, *g*^3^) ≥ *d*_*H*_(*g*^1^, *g*^2^) always holds. The equality means that no other local maximum lies “in between”.

## Notes

### Competing Interest Statement

The authors have declared no competing interest.

